# SpatialCorr: Identifying Gene Sets with Spatially Varying Correlation Structure

**DOI:** 10.1101/2022.02.04.479191

**Authors:** Matthew N. Bernstein, Zijian Ni, Aman Prasad, Jared Brown, Chitrasen Mohanty, Ron Stewart, Michael A. Newton, Christina Kendziorski

**Affiliations:** Morgridge Institute for Research, Madison, WI, 53715, USA; Department of Statistics, University of Wisconsin - Madison, Madison, WI, 53706, USA; Department of Dermatology, University of Wisconsin-Madison, Madison, WI, 53715, USA; Department of Biostatistics and Medical Informatics, University of Wisconsin - Madison, Madison, WI, 53792, USA

## Abstract

Recent advances in spatially resolved transcriptomics technologies enable both the measurement of genome-wide gene expression profiles and their mapping to spatial locations within a tissue. A first step in spatial transcriptomics data analysis is identifying genes with expression that varies spatially, and robust statistical methods exist to address this challenge. While useful, these methods do not detect spatial changes in the coordinated expression within a group of genes. To this end, we present SpatialCorr, a method for identifying sets of genes with spatially varying correlation structure. Given a collection of gene sets pre-defined by a user, SpatialCorr tests for spatially induced differences in the correlation of each gene set within tissue regions, as well as between and among regions. An application to cutaneous squamous cell carcinoma demonstrates the power of the approach for revealing biological insights not identified using existing methods.

## Introduction

Spatial transcriptomics (ST) experiments provide spatially localized measurements of genomewide gene expression, allowing investigators to address scientific questions that were elusive just a few years ago. Unlike single-cell RNA-seq (scRNA-seq) approaches in which a tissue sample is dissociated to produce a suspension of single cells thereby losing information about each cell’s location within the tissue, ST experiments retain spatial information and are therefore essential for comprehensively addressing questions associated with cell state and function when cell position and neighbors are crucial. By allowing investigators to address questions related to how adjacent cells communicate, how functional specialization is influenced by spatial location within a tissue, and how proximity among varying cell types affects downstream phenotypes, the ST technology has already enabled key insights into embryonic development (Srivatsan et al., 2021), nephrology (Ferreira et al., 2021), wound healing (Foster et al., 2021), brain function (Maynard et al., 2021), and cancer (Hunter et al., 2021; Ji et al., 2020; Moncada et al., 2020).

A first step in ST data analysis is identifying genes with expression that varies spatially, so-called spatially variable (SV) genes, and robust statistical methods exist to address this challenge (Andersson and Lundeberg, 2021; Edsgärd et al., 2018; Li et al., 2021; Sun et al., 2020; Svensson et al., 2018). While useful, SV genes alone do not fully describe, and in many cases cannot capture, important signals present in ST data. Specifically, cellular phenotypes are largely determined, and downstream phenotypes are largely affected, by spatially coordinated regulation (and deregulation, or reregulation) of expression among a group of genes. Canonical examples are found in cancer, where coordinated gene expression in immune cells changes based on spatial proximity to cancer cells, substantially affecting tumor progression and response to treatment (Demaria et al., 2019; Maier et al., 2020; Watson et al., 2021) and in spinal cord samples from ALS patients that show disease-relevant regional differences in coordinated expression among genes in microglia and astrocyte populations (Maniatis et al., 2019). These and numerous other studies (Hashimshony et al., 2015; Huisman et al., 2017; Li et al., 2016) demonstrate that spatial location can have a substantial effect on coordinated gene expression both within and among cell types, and that changes in coordinated expression have important implications for basic biology and medicine.

While SV methods provide information on changes in *average* expression, they do not specify how, or even if, genes within a group manifest *coordinated* expression. A group of SV genes may be independent, or dependent; or some subset may be dependent. In addition to identifying SV genes, knowledge of the dependence structure among a group of genes, and how it changes within and across tissue regions, is required to comprehensively describe expression dynamics underlying cell states in ST experiments.

Toward this end, we introduce SpatialCorr, a semiparametric approach for identifying spatial changes in the correlation structure of a group of genes. An overview is provided in Figure 1. Given a collection of gene sets and tissue regions pre-defined by a user (Figure 1a), SpatialCorr tests for spatially-induced differences in the correlation structure of each gene set within tissue regions, as well as between regions. Specifically, for a pre-defined set of genes, SpatialCorr estimates spot-specific correlation matrices using a Gaussian kernel (Figure 1b); region-specific correlations are estimated using all spots in a region (Figure 1d). SpatialCorr tests for spatially varying correlation <u>w</u>ithin each tissue <u>r</u>egion (the WR-test) using a multivariate normal (MVN) likelihood ratio test statistic that compares the MVN with spot-specific correlation estimates to an MVN with constant correlation estimated from all spots in the region (Figure 1c, f), and statistical significance is assessed nonparametrically. Specifically, p-values are computed via sequential Monte Carlo permutation (Besag and Clifford, 1991) and FDR is controlled across regions using Benjamini-Hochberg (Benjamini and Hochberg, 1995). For each set on test, region-specific p-values are provided to highlight spatial locations showing the most significant internal variation (Figure 1c) and gene-pairs within the group are also ranked by pairwise p-values so that those pairs contributing most to the differential correlation (DC) call can be easily identified. The test for identifying correlations that change between regions (the BR-test) is not well defined for regions showing internal changes in correlation. Given this, we only consider those regions for which the correlation structure among genes shows no significant change according to the WR-test. Then, for each pair of regions, differences in correlations among the genes are identified using the between-regions (BR) test (Figures 1e, f). As with the WR-test, FDR-adjusted p-values for the BR-test are computed via permutation.

**Figure 1.**
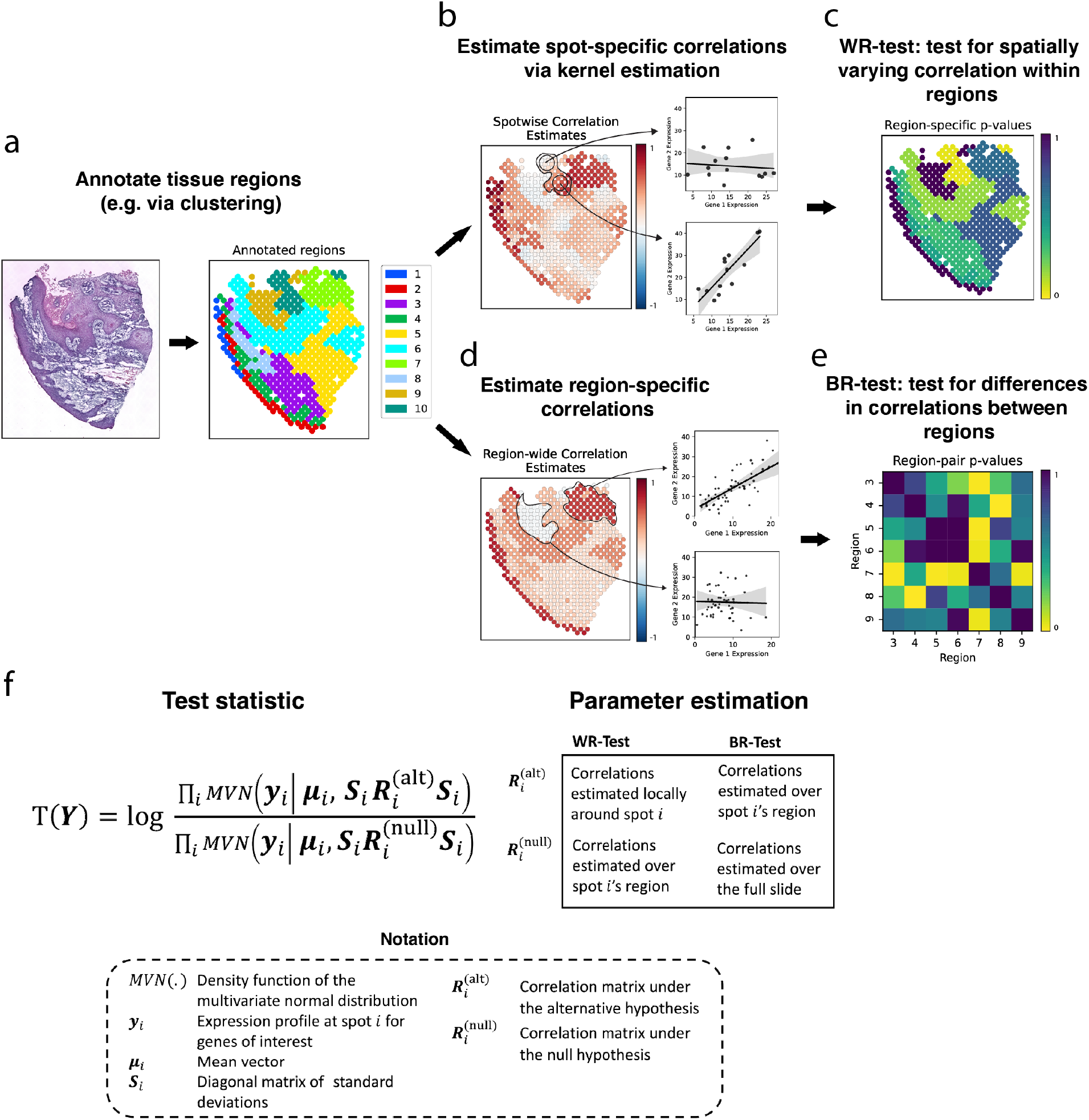
SpatialCorr overview. A schematic workflow of the analyses performed by SpatialCorr. (**a**) First, distinct tissue regions are identified (e.g. via clustering or manual annotation). (**b**) For a pre-defined set of genes, since spots do not contain replicates, SpatialCorr estimates correlations at each spot using a Gaussian kernel. In this example, regions 1 and 2 have too few neighbors for the Gaussian kernel and, consequently, were removed by the effectiveneighbors filter. (**c**) SpatialCorr tests for spatially varying correlation <u>w</u>ithin each tissue <u>r</u>egion (the WR-test); p-values are estimated via permutation with FDR controlled across regions using Benjamini-Hochberg. The WR-test identifies region 10 as significant. (**d**) Regions without spatially varying correlation are identified (here, regions 3-9) and region-specific correlations are estimated using all spots in those regions. (**e**) The pre-defined set of genes is tested for differences in correlation structure between each pair of regions using the BR-test. As in the WR-test, p-values for the BR-test are computed via permutation and FDR is controlled across pairs using Benjamini-Hochberg. (**f**) An overview of the MVN likelihood ratio test statistic used by SpatialCorr. The WR test compares spot-specific correlations with correlation calculated across the region; the BR-test compares region-specific correlations with correlation calculated across the pair of regions being compared (when testing for differences between two regions) or across the entire slide (when testing for differences among regions across the entire slide).

SpatialCorr relies on sequential Monte Carlo (SMC) permutation (Besag and Clifford, 1991) to efficiently compute p-values, and is implemented in an easy-to-use Python package. Consequently, it is computationally efficient and scalable to hundreds of gene groups; it also integrates directly into existing spatial transcriptomics analysis pipelines implemented in Scanpy (Wolf et al., 2018). Finally, the SpatialCorr software contains the first approach for simulating realistic count data with spatially varying correlation structure. In addition to evaluating SpatialCorr, the simulation framework developed here is expected to prove useful in benchmarking studies as additional DC methods for spatial experiments are developed.

In contrast to scHOT (Ghazanfar et al., 2020), an approach for identifying changes in a gene’s variance or in gene-gene pairwise correlations across the entire tissue, SpatialCorr considers the full dependence structure among a group of genes, as represented by the gene groups’ correlation matrix. In addition, SpatialCorr conducts DC tests within a tissue region, or between tissue regions, resulting in improved localization of the DC signal. SpatialCorr also directly accounts for differences in latent expression levels across distinct tissue regions, avoiding false positives due to mean expression that varies across regions. Since spurious correlation can be induced by a latent factor, it is important to directly model spatial changes in mean expression to avoid false positives in DC tests (Supplementary Note S1).

## Results

### SpatialCorr increases the power to identify groups of genes having spatially varying correlation within and between tissue regions

We conducted several simulations to evaluate the performance of SpatialCorr under a variety of conditions. While methods are available to simulate Gaussian data having a predefined correlation, these methods are not easily extended to count data with correlation that varies spatially (Inouye et al., 2017). To simulate realistic ST data with spatially varying dependencies between genes, we developed a novel simulation framework based on the Poisson-lognormal model (Figure 2a). Given a spatially varying covariance matrix, the framework provides spatially resolved UMI count data having marginal distributions and spot-specific size factors that match input experimental data while controlling the correlation between a group of genes at each spot. Pre-specifying a spatially varying covariance matrix is challenging given the requirements that it be positive semi-definite. To address this challenge, the simulation framework within SpatialCorr provides a function to generate smoothly varying covariance matrices using a Gaussian process model.

**Figure 2.**
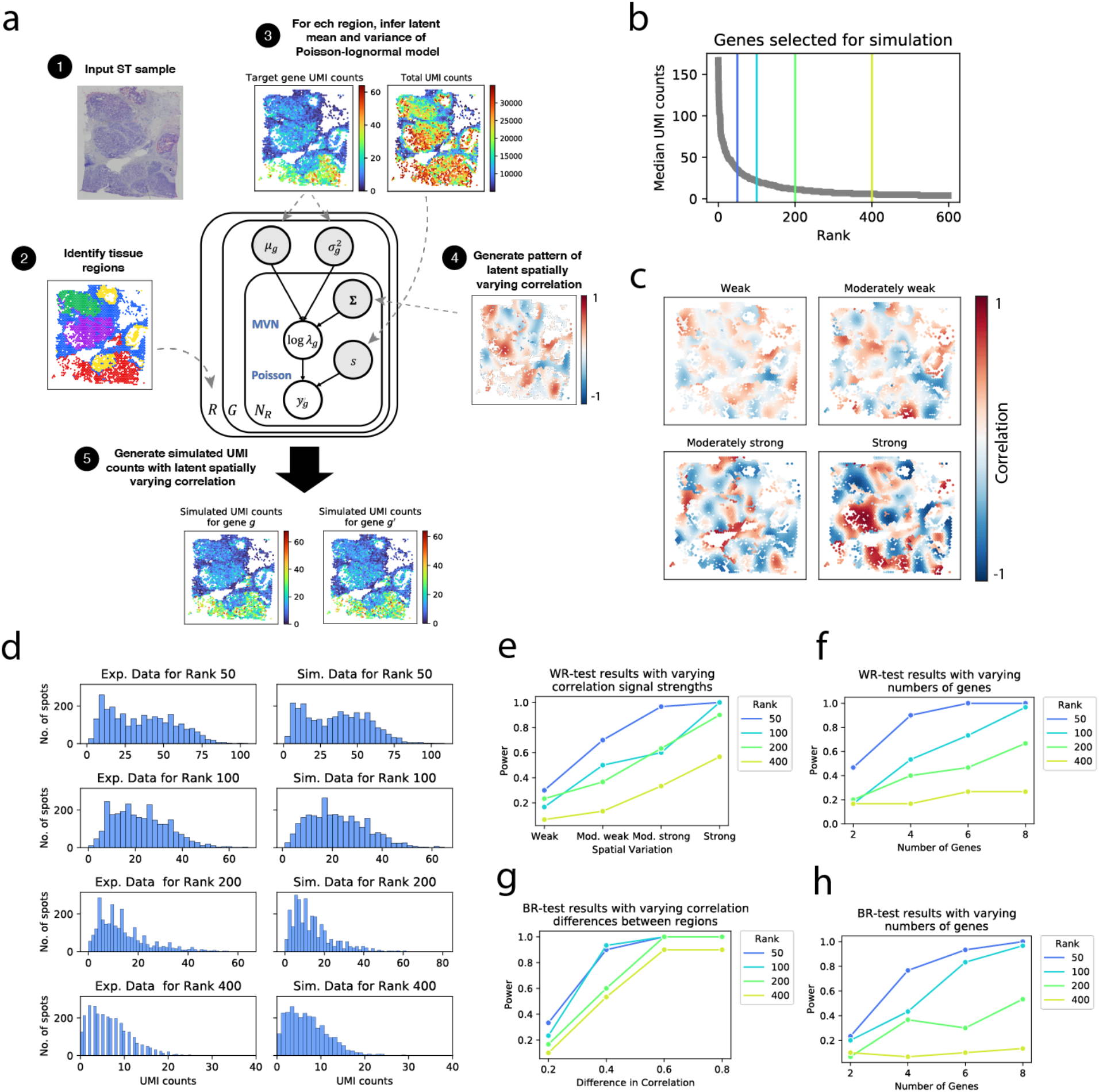
Simulation framework and results. An overview of the simulation framework is shown in **(a)**. Given an input ST sample, tissue regions are identified (e.g. via clustering or manual annotation). A user then specifies the number of genes to simulate and selects a corresponding set of target genes in the spatial transcriptomics sample upon which the simulations are based. For each target gene, the latent parameters in the Poisson-lognormal model are inferred on a per-region basis. A pattern of spatially varying latent correlation is then generated using a Gaussian process sampling procedure that is tuned to match the correlations observed in experimental data. Finally, counts are generated from the Poisson-lognormal model. **(b)** In an effort to ensure that results provide relevant information, data is simulated at realistic gene expression levels. Specifically, genes are rank ordered by total UMI counts. Here, four genes were selected based on expression (rank 50, 100, 200, and 400); the 400th ranked gene is considered lowly expressed, rank 200 moderately low, rank 100 moderately high, and rank 50 highly expressed. Four correlation levels (low, moderately low, moderately high, high) were also considered based on those observed in real data. (**c**) Heatmaps depicting four examples of the latent spatially varying correlation patterns generated for each of the four correlation levels. **(d)** Histograms comparing the distribution of UMI counts in the experimental data (left) to the simulated data (right) for each of the four expression levels. (**e**) The average power of the WR-test is shown for varying correlation and expression levels (averages taken over 30 simulated datasets). (**f**) The average power of the WR-test is shown for varying correlation and expression levels of gene sets of size 2, 4, 6, and 8 (averages taken over 30 simulated datasets). (**g, h**) Similar to (e) and (f), but for the BR-test. Instead of varying the amount of underlying correlation among genes, we vary the difference in correlation between regions.

We carried out four sets of simulations, Sims I-IV, using as input a breast cancer sample assayed via the 10x Visium platform. To ensure that the simulations provide both realistic and comprehensive results, genes from the input dataset were rank ordered by median UMI counts and four genes were selected as shown in Figure 2b. Data was simulated for genes having low expression (matching the 400th ranked gene), moderately low (200th ranked gene), moderately high (100th ranked gene), and high (50th ranked gene). Four correlation levels (weak, moderately weak, moderately strong, strong) were also considered based on those observed in real data (Figure 2c, Supplementary Note S2). In Sim I, we simulate a pair of lowly expressed genes at each of the four correlation levels (30 replicates per level); the simulation was repeated for gene pairs having expression in the moderately low, moderately high, and high expression ranges. The marginal distributions for simulated genes at each of the expression levels are shown in Figure 2d. Sim II is similar, but considers groups of genes of size 4, 6, and 8.

Results demonstrate the increases in power observed with increasing expression and correlation levels. Specifically, Figure 2e shows that the WR-test in SpatialCorr has low power to identify weakly correlated gene pairs, even those that are highly expressed. However, there is reasonable power to identify gene pairs with stronger levels of correlation. Figure 2f shows that SpatialCorr’s power increases substantially when the correlation structure across multiple genes is considered jointly. When groups of 4 genes having weak correlation are considered, for example, power is increased from 47 to 90%; the power is further increased to nearly 100% when 6 genes are considered. False-positive rates are well-controlled across simulation settings; for each expression level, we simulated pairs of genes from the null scenario in which there is no spatially varying correlation in each region (200 samples for each expression level). The false positive rate (FPR) across all null samples was 0.034 when using a p-value cutoff of 0.05 to make a positive call.

Sim III evaluates the BR-test. Specifically, we simulate a pair of lowly expressed genes with correlation that differs by 0.2 between regions but remains constant across spots within each region; the simulation was repeated for gene pairs having expression in the moderately low, moderately high, and high expression ranges. For each expression level, the simulation was repeated with increasing differences in correlation of 0.4, 0.6, and 0.8 between regions. Sim IV is similar, but considers groups of genes of size 4, 6, and 8.

Results demonstrate the increases in power observed with increasing expression and difference in correlation between regions. Specifically, Figure 2g shows that the BR-test in SpatialCorr has low power to identify correlated gene pairs whose correlation differs minimally between regions, even those that are highly expressed (Figure 2g). However, the power increases significantly when the difference in correlation between regions increases to 0.4 and above. Figure 2h shows that SpatialCorr’s power also increases substantially when the correlation structure across multiple genes is considered jointly. When groups of 4 genes with pairwise correlations that differ by 0.2 between regions are considered, for example, power is increased from 23 to 77%; the power is further increased to 93% when 6 genes are considered. As with the WR-test, the false-positive rate is well controlled (at 0.04 when using a p-value cutoff of 0.05 to make a positive call).

### SpatialCorr reveals spatially varying correlation among genes within and between regions of cutaneous squamous cell carcinoma

To further assess the performance of SpatialCorr, we applied it to clinical ST data from human cutaneous squamous cell carcinoma (cSCC), the second most common cancer worldwide (Ji et al., 2020; Rubió-Casadevall et al., 2016). In spite of its prevalence, the molecular basis of cSCC remains poorly understood and surgery is the main treatment (Dotto and Rustgi, 2016). ST experiments analyzed with SpatialCorr offer the chance to reveal new therapeutic insight beyond traditional ST analysis, including previously unknown regulatory networks. In Ji *et al*., ST data was obtained from four patients. Three of the four patients (patients 2, 4, and 6) had the histopathologic subtype of well-differentiated cSCC whilst patient 10 had a more aggressive histopathologic subtype of moderately differentiated cSCC (Rowe et al., 1992). SpatialCorr was applied to each dataset to identify spatially varying correlation among groups of genes within or between regions. We considered 836 groups of genes defined by GO categories involved in immune response, cell adhesion, proliferation, and others relevant to tumorigenesis (Dotto and Rustgi, 2016).

Results from the WR and BR-tests are summarized in Supplementary Tables S1-8. Consistent with the nature of the cSCC tumors samples, the top GO term identified by either the WR or BR-tests in all four patient samples was “keratinocyte differentiation”. This GO term is enriched for keratin genes, which are primarily structural proteins of keratinocytes, but which have emerging roles in tumor invasiveness and metastasis (Karantza, 2011). Keratins are obligate heterodimers in which type 1 keratins (e.g. KRT1, KRT5, and KRT6) classically bind to type 2 keratins (e.g. KRT10, KRT14, KRT16, and KRT17) and are therefore expected to be highly correlated in skin (Moll et al., 2008). This presents a convenient test case for SpatialCorr and we indeed identified strong correlation between expected type 1 and type 2 pairs such as KRT1 and KRT10, as well as KRT5 and KRT14, throughout the epidermis and much of the tumor (Figures 3a-c; Figure SF1). Less well understood are changes in the strength of typical keratin correlation patterns, as well non-canonical pairings, that have been observed in skin disease (Quigley et al., 2016; Toivola et al., 2015). To examine the spatially varying patterns among these keratins more closely, we identified subgroups of keratins having similar correlation patterns by clustering the pairwise correlations estimated from SpatialCorr. Figure 3f shows five subgroups derived from the moderately differentiated cSCC sample. A focus on one of the subgroups shows strong positive correlation between keratin pairs in the tumor region that decreases in the non-tumor region; the strongest pairs in the tumor region include KRT17 and KRT16 as well as KRT17 and KRT5. Most pairwise correlations in this subgroup are highest in the tumor region defined by cluster 7 (Figure 3b; Figure 3h) reflecting coordinated regulation of these keratins within the region. This region corresponds to a novel cell subpopulation unique to cSCC made up of tumorspecific keratinocytes (TSKs) that was identified in Ji *et al*. 2020 as putatively involved in tumor progression, immunosuppression, and tumor invasiveness. While KRT17 and KRT16 have been independently associated with proliferation (Paramio et al., 1999) and aggressiveness in a number of cancers (Han et al., 2021; Huang et al., 2019; Liu et al., 2020; Nair et al., 2021), their coordinated regulation has not been studied. The strong correlation identified by SpatialCorr suggests an increased level of coordinated regulation within the TSK region, providing further insights into this novel subpopulation.

**Figure 3.**
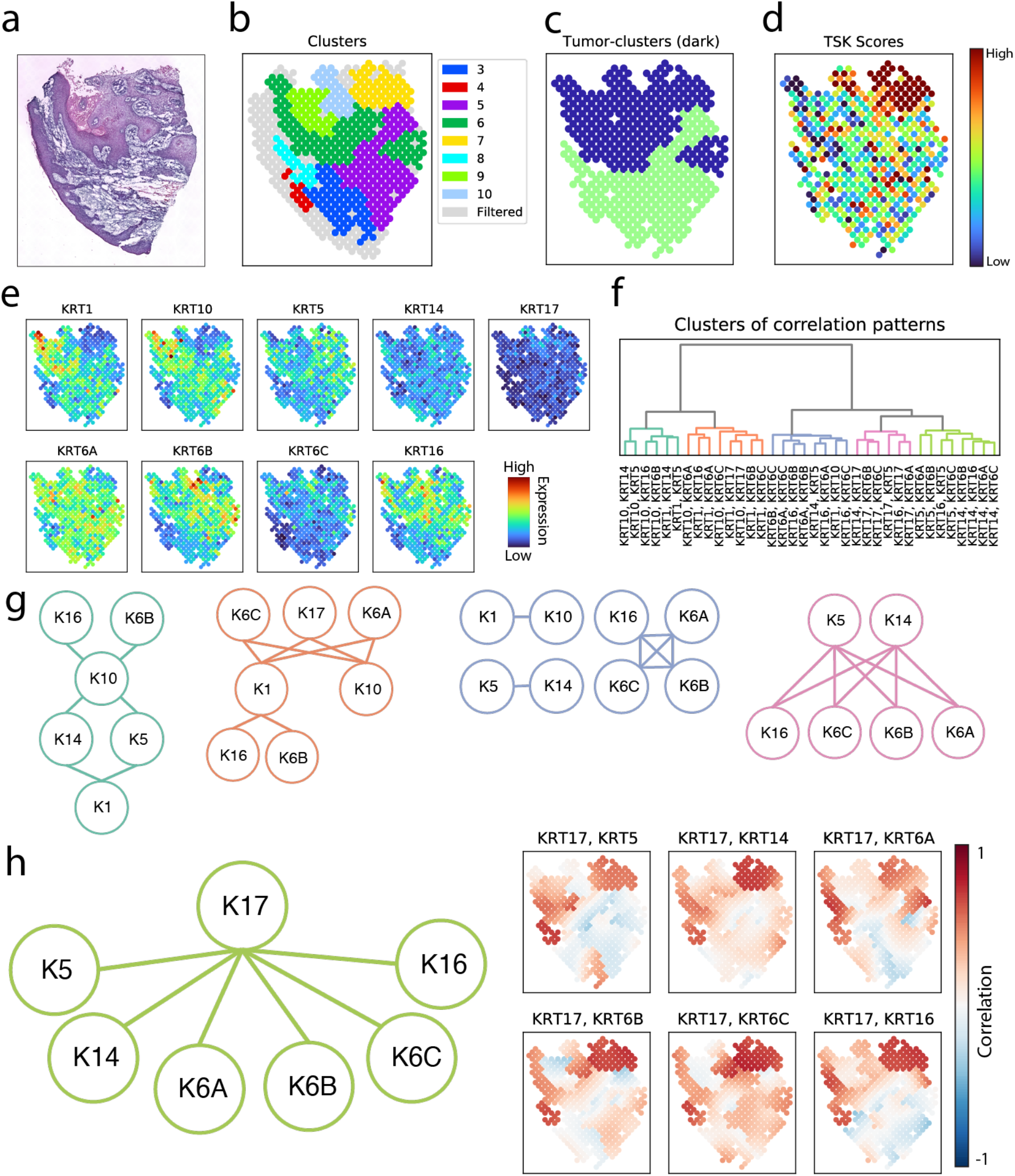
Analysis of cutaneous squamous cell carcinoma data. (**a**) H&E stain of moderately differentiated cSCC tumor (patient 10) from Ji *et al*. (2020). (**b**) Spots colored by their assigned BayesSpace clusters; spots that were removed by the effective-neighbors filter (having too few neighbors to estimate a correlation via the Gaussian kernel) are colored in grey. Here, all spots from clusters 1 and 2 were removed by the filter. (**c**) Spots colored according to whether they belong to a cluster that is mostly tumor. (**d**) Expression of 9 keratin genes in the keratinocyte differentiation GO category that is identified by both the WR and BR-test in SpatialCorr as having spatially varying correlations. (**e**) Spots colored according to their tumor-specific keratinocyte (TSK) score, which measures the enrichment of the tumor-specific keratinocyte population discovered by Ji *et al*. (2020). (**f**) Spot-specific kernel estimates of correlation between all pairs of keratin genes were calculated and the pairs of genes were clustered according to their correlation pattern across the slide. **(g)** Subgroups were identified by cutting the dendogram shown in (f). Each subgroup is shown as a graph consisting of genes where an edge is drawn between genes if the corresponding gene-pair belongs to the subgroup and each graph is colored according to (f). **(h)** A subgroup involving KRT17 is highlighted (left); kernel estimates of correlation between each pair in the subgroup (right). We note high correlation between KRT17 and genes KRT5, KRT14, KRT6, and KRT16 in the region of the tumor enriched for the TSK population.

### SpatialCorr reveals temporally varying correlation among genes within differentiating hematopoietic cells

While we have focused on ST data, SpatialCorr can be applied to any gene expression dataset for which a distance metric is defined between samples. To demonstrate, we consider a study of hematopoiesis from (Paul et al., 2015) where scRNA-seq data is profiled in differentiating mouse myeloid and erythroid hematopoietic cells (Figure 4a); distance between cells is defined by the shortest path distance along the nearest-neighbors graph (Figure 4b). SpatialCorr was applied to test for temporally varying correlation within 227 groups of genes defined by GO terms related to myeloid and erythroid cell function and differentiation. The top gene set identified was Antimicrobial Humoral Response, which includes the genes Ctsg, Elane, and Prtn3. SpatialCorr finds increased pairwise correlation along the monocyte and neutrophil trajectories, with lower correlation along the other branches (Figure 4c). These genes are thought to be regulated by a common promoter and were recently discovered to jointly catalyze histone H3 amino terminus proteolytic cleavage in a process that controls hematopoietic cell differentiation (Cheung et al., 2021).

**Figure 4.**
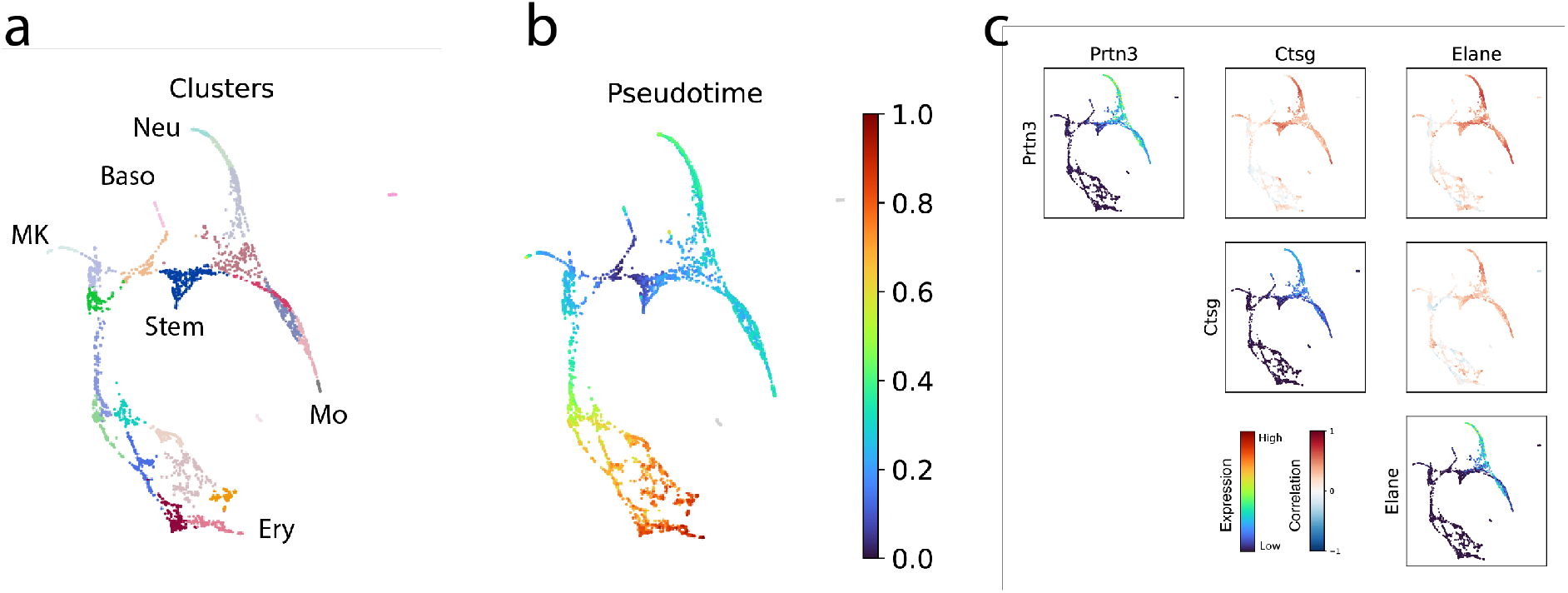
Analysis of differentiation hematopoietic cells. (**a**) Force-directed layout of the nearest-neighbors graph of cells colored by Louvain clusters (Blondel et al., 2008). Cells are differentiating from hematopoietic stem cells (Stem) into neutrophils (Neu), monocytes (Mo), basophils (Baso), megakaryocytes (MK), and erythrocytes (Ery). (**b**) Force-directed layout of cells colored according to their estimated pseudotime. (**c**) Correlation analysis for the genes Prtn3, Elane, and Ctsg. Plots along the diagonal depict each gene’s expression. Off-diagonal plots depict the cell-wise kernel estimates of the correlation between the gene pair. We note high correlation between these genes along the neutrophil and monocyte trajectories.

### The SpatialCorr package is an efficient and comprehensive suite of tools for analyzing and visualizing spatial correlation patterns

SpatialCorr is implemented in a fast and user-friendly Python package that implements a variety of functions for studying spatially varying correlation structure. In addition to the WR and BR-tests that facilitate identification of gene sets having significantly varying correlation within or between regions, additional functions within SpatialCorr enable a user to compute and visualize per-spot kernel estimates of correlation (as shown in Figure 3h, right panel), confidence intervals around these estimates (Figure SF2a), regional estimates of correlation (Figure SF2b), and clusters of correlation patterns within groups of gene pairs (Figure 3f). For further downstream analyses, the package can be easily integrated into existing analysis pipelines implemented with Scanpy. In addition to these analysis capabilities, SpatialCorr includes a simulation framework for simulating ST data which we expect will prove useful in the development and evaluation of future methods.

SpatialCorr utilizes Python’s efficient NumPy library (Harris et al., 2020) which provides relatively fast performance. When computing an SMC p-value, permutations are computed sequentially and terminated early if the p-value is deemed to be high. In this way, fewer permutations are generated when the p-value is high thereby diverting computational resources towards those gene sets that produce low p-values, which require precise estimates for downstream methods that account for multiple testing. To evaluate timing on analysis of pairs, we considered 20 highly expressed gene pairs from a large sample of brain tissue consisting of 4,226 spots from (Maynard et al., 2021). The WR-test takes approximately 2.8 minutes per pair to estimate SMC p-values using 1000 permutations on a single CPU core; the BR-test takes approximately 3.8 minutes per pair (17 and 30 minutes per pair are required, respectively, to estimate exact p-values). In contrast, scHOT takes over 9 hours per pair running 1,000 permutations on one CPU core, indicating that scHOT does not easily scale to analyses that involve multiple gene pairs in ST samples with many spots. Finally, to further improve speed, the SpatialCorr permutation test can be parallelized to utilize multiple CPU cores.

## Discussion

A fundamental task in ST experiments is identifying genes with expression levels that vary across a tissue sample, and a number of differential expression (DE) methods have been developed toward this end (Andersson and Lundeberg, 2021; Edsgärd et al., 2018; Li et al., 2021; Sun et al., 2020; Svensson et al., 2018). Although a critical first step, DE measures do not capture many important types of differential regulation. Specifically, complex phenotypes may arise from a de- or re-regulation that does not significantly affect a gene’s *average* expression level, but rather affects the ways in which a gene’s expression changes with respect to other genes. SpatialCorr provides the first approach for identifying groups of genes with spatially varying correlation within or between tissue regions. We expect it will be a useful complement to existing ST analysis methods.

## Methods

Test statistics in SpatialCorr’s WR-test and BR-test are derived from normal distribution theory, although the permutation analysis assures validity beyond the normal data model. Specifically, let ***Y*** be an *n* × *m* normalized expression matrix for a set of *m* genes with expression profiled at *n* spots. The alternative and null models are then

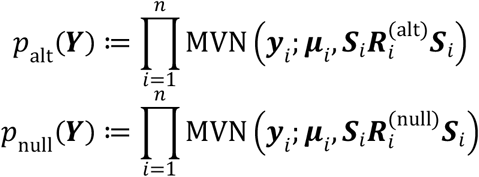

respectively, where ***y**_i_* is the *m*-dimensional vector of gene expression values at spot *i*, ***μ**_i_* is the *m*-dimensional mean expression vector for the set of genes, ***S**_i_* is a diagonal matrix consisting of the genes’ standard deviations, and 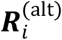 and 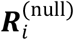 are the *m* × *m* correlation matrices at spot *i* under the alternative and null models, respectively. The test statistic, *T*(***Y***), is the log-likelihood ratio between these models. The means of both models are allowed to vary spot-to-spot to ensure that the test is testing for spatial differences in correlation rather than spatial differences in mean expression. The mean of each gene is estimated using kernel estimation (Yin et al., 2010):

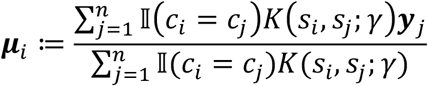

where *s_i_* and *s_j_* are the coordinates for spots *i* and *j* respectively, *c_i_* and *c_j_* are the tissue region id’s for spots *i* and *j* respectively, 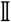 is the identity function, *K* is the kernel function, and *γ* is the bandwith parameter. SpatialCorr uses a radial basis kernel for *K*. We chose a default bandwidth parameter of five when operating on spot-coordinates from the 10x Visium platform as we have found this bandwidth parameter small enough to detect granular changes in spatial correlation.

### WR-test

Under the null model, the diagonal entries of the covariance matrix, 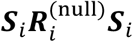, are allowed to vary spot-to-spot to ensure that the test is testing for spatial differences in correlation rather than for spatial differences in variance. Like the means, the variance of each gene *g* is estimated using kernel estimation:

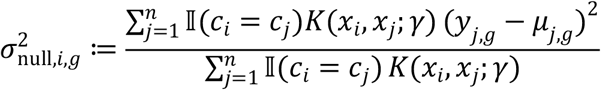

Under the null model, the off-diagonal entries of the covariance matrix are calculated as follows: First, for each tissue region *C*, the Pearson correlation matrix among the set of genes, ***R**_C_*, is estimated using the data for all spots in that region. Then, for each spot, the off-diagonal entries of the covariance matrix – that is, the covariance estimates between genes *g* and *g′* – are calculated as 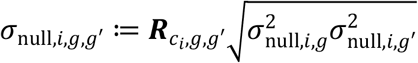 where *c_i_* is the region in which spot *i* belongs and ***R**_c_i_,g,g′_* is the correlation between genes *g* and *g′* in region *c_i_*.

For the alternative model, the entire covariance matrix, including the off-diagonal entries, is estimated via kernel estimation (Yin et al., 2010):

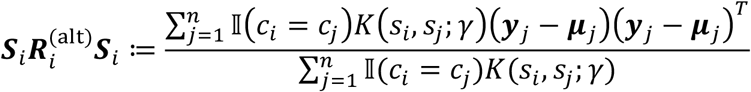

### BR-test

Just as in the WR-test, the null and alternative models differ according to off-diagonal entries of the covariance matrix. In the alternative model, the off-diagonal entry representing the covariance between two genes *g* and *g′* is given by 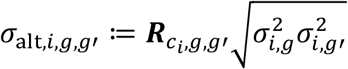 where *c_i_* is the tissue region of spot *i* and ***R**_c_i_,g,g′_* is the Pearson correlation estimated in region *c_i_*. Under the null model, the Pearson correlation is calculated between the two genes using all the spots as 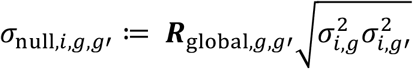 where ***R**_global,g,g′_* is the Pearson correlation estimated globally using all of the spots.

### Filtering spots

Estimates of spot-specific parameters used in SpatialCorr’s test statistic will have high variance for spots with few neighbors due to a small number of spots contributing disproportionately to the estimates. For this reason, SpatialCorr filters spots with a small number of neighboring spots by setting a filter on the number of “effective neighbors” -- that is, the effective sample size used in the calculation. For spot *i*, effective_neighbors_*i*_ 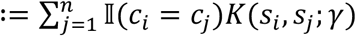. The default effective-neighbors threshold is set to 10.

### Assessing statistical significance

We calculate the significance of *T*(***Y***) via a permutation test so that the false positive rate is robustly controlled. The null hypothesis in the WR-test specifies that there is no spatially varying correlation within each region and thus spots (i.e., normalized expression profiles) are exchangeable over this region and may be permuted without affecting their joint null probability density. For the BR-test, each tissue regions’ spots are first zero-mean centered and then the residual expression profiles are permuted between spatial coordinates across the entire slide (or across the pair of regions if we are testing for differing correlation structure between a region-pair).

In order to speed up the computation, SpatialCorr computes sequential Monte Carlo (SMC) p-values. Behind the idea of SMC p-values is the insight that one does not require as precise an estimate of a large p-value as one does for a small p-value. That is, if a p-value is large, we need few samples from the null distribution to safely conclude that the null hypothesis cannot be rejected. In contrast, one requires precise estimates of small p-values in order to perform more precise downstream analyses and correction for multiple hypothesis testing. To compute an SMC p-value, one sets a predefined threshold *t* (by default, we set *t* = 20) and generates samples from the null distribution by calculating the test statistic *T*(***Y***) on sequentially generated, random permutations until *t* samples from the null exceed *T*(***Y***). The p-value is then computed as *p* ≔ *t/l* where *l* is the total number of permutations. For extremely small p-values that may require an exorbitant number of permutations before termination, the algorithm is terminated at a predefined *g* − 1 permutations (by default, we set *g* = 10,000), and estimates the p-value as *p* ≔ (*u* + 1)/*g* where *u* is the number of null samples that exceed *T*(***Y***).

### Computing confidence intervals around correlation estimates

We calculate a confidence interval around the spot-specific kernel estimates of the correlation at each spot using a non-parametric bootstrap. Specifically, a fixed radius is considered around each spot and a set of bootstrap samples of spots are taken from this neighborhood. For each bootstrap sample, the kernel estimate of the correlation is calculated as described above; and a confidence interval is computed using the bootstrapped correlation estimates.

### Simulation framework

Our goal is to simulate realistic ST data where the covariance for select sets of genes varies smoothly across the slide. Toward this end, each simulated dataset requires a case study input dataset from which parameters are estimated. The simulation framework is built upon two models. The first model generates smoothly varying latent correlations between genes at each spot. The second model is a lognormal-Poisson model that uses these correlations as parameters to generate UMI counts.

For spot *i*, counts for two genes are simulated as Poisson: *y*_*i*,1_ ~ Poisson(*s_i_λ*_1_) and *y*_*i*,2_ ~ Poisson(*s_i_λ*_2_) with log*λ*_1_, 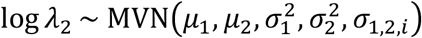. Here, *N* is the total number of spots, *s_i_* is the total UMI count at spot *i*, and *y_i,j_* is the UMI count for gene *j* at spot *i*. Synthetic measurements are mutually independent among spots; the dependencies arise between genes within spots. The means, *μ*_1_ and *μ*_2_, and variances, 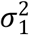 and 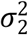, are estimated using the experimental data. Specifically, for each gene, we use the posterior means from a simple hierarchical Bayesian model that uses a normal prior for the means and a truncated Cauchy prior for the variances. When simulating UMI counts from distinct tissue regions, we perform the aforementioned process on each region independently using different parameters (based on different pairs of genes) for each region.

The covariance parameters, *σ*_1,2,*i*_, across all spots *i* ∈ [*N*], are generated according to a Gaussian-process model. First, we compute the kernel matrix between spots using a radial basis function kernel 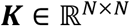 where each element is computed as

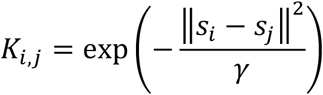

where *s_i_* and *s_j_* are the spot-coordinates for spot *i* and spot *j*, and *γ* is the bandwidth parameter that sets the width of the radial basis function. Given the kernel, latent Fisher-transformed correlations are sampled from a Gaussian process atanh (***r***) ~ MVN(**0**, *c**K***).

Note that *c* is referred to as the “covariance strength” parameter which increases or decreases the variance of the correlations across the slide. A high value for *c* will produce stronger correlations (more correlation values close to −1 and 1) whereas small values for *c* will produce weaker correlations (closer to 0). Fisher-transformed correlations are converted to Pearson correlations and the covariance between the two genes is then defined as 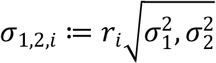 where *r_i_* is the Pearson correlation between the two genes at spot *i* (i.e., the *i*th element of ***r***). Note that 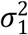 and 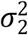 are estimated from the experimental data as previously described.

To simulate UMI counts for sets of genes with spatially varying correlation, we use the same Poisson-lognormal-based model as was described previously for simulating pairs of genes. However, we augment the process for generating smoothly varying latent correlations due to the requirement that the covariance matrix must be positive semidefinite. For each gene *g* in the gene set, we generate a normally distributed vector of length *N*, ***v**_g_* ~ MVN(**0**, *c**K***), where again 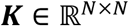 is a kernel matrix, and *c* controls the variability of the correlation values across spots. For each spot *i*, we simulate latent expression profiles for all genes by sampling from a MVN distribution 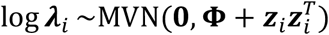 where **Φ** is a diagonal matrix that stores the variance of each simulated gene and ***z**_i_* ≔ [*v*_1,*i*_ *v*_2,*i*_ … *v_G,i_*]^*T*^ where *G* is the total number of genes in the target gene set. In short, for a given gene *g*, the random variables *v_g,i_* vary smoothly across the spatial coordinates and thus, for a given pair of genes, *g*_1_ and *g*_2_, the value *v*_*g*_1_,*i*_*v*_*g*_2_,*i*_ (i.e., an off-diagonal entry of 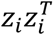) varies smoothly. In this way, we can randomly generate a positive semidefinite matrix at each spot such that they vary smoothly, elementwise, across the spatial coordinates while controlling the resolution of these changes (Hoff and Niu, 2012). Note that the covariance strength parameter, *c*, scales the variability in the entries of the 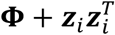 matrices.

### Simulated data

Simulated data was generated based on Sample *V1_Breast_Cancer_Block_A_Section_1*, which was downloaded from 10x Genomics using the Scanpy Python library via the *visium_sge* function. This sample is also available to download from 10x Genomics website: (https://www.10xgenomics.com). We generate simulated data under four simulation scenarios, described as follows.

#### SimI

Two genes and five tissue regions with spatially varying correlation within each region. We simulate datasets for gene pairs with four levels of expression corresponding to the 50^th^, 100^th^, 200^th^, and 400^th^ gene in the experimental data ranked according to their median UMI counts. The strength of the correlation, as set by the correlation strength parameter, *c*, is defined to be zero (*c* = 0.0), weak (*c* = 0.05), moderately weak (*c* = 0.1), moderately strong (*c* = 0.2), and strong (*c* = 0.4). These five parameter values for *c* were found to generate data with realistic spatially varying correlation (see Supplemental Note S2).

#### SimII

Multiple genes and five tissue regions with spatially varying correlation within each region. We simulate datasets for genes with four levels of expression as per SimI. A total of nine genes were simulated using a covariance strength parameter of *c* = 0.1. We note that the effect of *c* on the variability of generated spot-wise correlations depends on the number of genes being simulated. We found a value of *c* = 0.1 in the nine-gene setting generated a similar strength of correlation as was produced by *c* = 0.05 (the “weak” setting) in the two-gene scenario of SimI. For an example of simulated pairwise correlation patterns for four genes, see Figure SF3a.

#### SimIII

Two genes and five tissue regions with no varying correlation within each region but differing correlation between regions. We simulate datasets for gene-pairs with four levels of expression as per SimI. We grouped the regions into two groups where each group shared the same internal correlation, but the correlation between the groups of regions differed. Differences in the correlation between the regions was set to zero, 0.2, 0.4, and 0.8. For an example of simulated pairwise correlation patterns, see Figure SF3b.

#### SimIV

Multiple genes and five tissue regions with no varying correlation within each region but differing correlation between regions. We simulate datasets for genes with four levels of expression as per SimI. A total of nine genes were simulated for each dataset. We grouped the regions into two groups where each group shared the same internal correlation structure, but the correlation between the groups of regions differed. Specifically, in one group, the correlation between all pairs of genes was zero, but in the second group, it was set to 0.2. For an example of simulated pairwise correlation patterns for four genes, see Figure SF3c.

### Calculation of TSK-scores in cSCC data

We used the TSK marker genes provided in the Supplementary materials of Ji *et al*. (2020). For each gene and each spot, we compute the z-score of the Dino-normalized gene expression values. We define the TSK-score at each spot to be the sum of z-scores across all TSK-markers.

### Normalization

It is often observed in spatial transcriptomics datasets that the total UMI counts detected at each spot (hereafter referred to as *library size*) vary significantly across the tissue slide. Moreover, this variation is not random as regions of high or low counts tend to cluster together. Moreover, because kernel estimation of the correlation is susceptible to changes in measured expression (see Supplemental Note 1), these changes in library size threaten to induce spurious correlation across the slide. For this reason, it is crucial to normalize the expression data to account for changes in library size across the slide. For all datasets used in this study, including simulations, we used the Dino algorithm to perform normalization because it was shown to better normalize for changes in library size than existing state-of-the-art methods (Brown et al., 2021).

### Publicly available case study datasets

For the analysis of the cSCC data from Ji *et al*. (2020), we used samples from patients 2, 4, 6 and 10 (GEO accessions GSM4284316, GSM4565823, GSM4565826 and GSM4284326, respectively). Clustering was performed using BayesSpace (Zhao et al., 2021) (v1.2.0 for R v4.1). We followed the procedure outlined in the vignette for the BayesSpace R package. The number of clusters for each sample was selected by choosing the elbow of the curve plotting the log-likelihood of the BayesSpace model versus the number of clusters as recommended by Zhao *et al*. Based on visual inspection, the cluster boundaries matched well with the leading edge of the tumor region as identified by Ji *et al*. (2020), and therefore, we considered a cluster to be tumor-enriched if its spots fell within the tumor region.

For the analysis of the hematopoiesis single cell data, cells were normalized, ordered by psuedotime, and annotated according to the Scanpy tutorial for trajectory inference (https://scanpy-tutorials.readthedocs.io/en/latest/paga-paul15.html). Specifically, normalization was performed using log counts per million followed by z-score normalization. Pseudotime was calculated using graph diffusion (Haghverdi et al., 2016). Cells were also normalized separately via Dino and the Dino-normalized expression values were used to run SpatialCorr. Lastly, we used the shortest path distances between cells along the four-nearest-neighbors graph and used these pairwise distances to compute the Guassian kernel used by SpatialCorr.

For the experiments that tested SpatialCorr’s execution time, we used sample 151507 from the SpatialLIBD project (Maynard et al., 2021). We used the author-provided “layer_guess” annotations of the cortical layers to designate the regions.

### Selection of gene sets for analysis

For each patient, we applied SpatialCorr to 836 groups of genes defined by GO categories known to be relevant to cancer, the skin, and skin development, such as those related to cellular proliferation, cell-cell adhesion, cell death, DNA repair, immune response, and others. GO terms were retrieved in batches by first determining a set of target terms (e.g. “DNA repair”) and for each term, the set of GO terms were identified that contained the given term as a substring. For each GO term, the top 15 most highly expressed genes (measured by the fraction of spots on the slide with at least one UMI count) were used within each category after filtering genes not detected in at least 0.2 of spots. GO categories having fewer than 5 selected genes were not used in order to obtain gene sets of roughly equal size. From the 836 gene sets, this resulted in consideration of 134 gene sets for patient 6, 287 gene sets for patient 10, 539 for patient 4, and 555 for patient 2. We curated a collection of 227 GO categories related to hematopoiesis using the same workflow used in the analysis, but starting with key terms related to immune cell function and hematopoiesis (e.g., “hematopoietic stem”). From the 227 gene sets, this resulted in consideration of 64 gene sets.

### Code availability

SpatialCorr is implemented as an open-source Python package and is available on GitHub (https://github.com/mbernste/spatialcorr). The software implementing the simulation framework is also available as an open-source Python package that is available on GitHub (https://github.com/mbernste/spatialcorr-sim). Lastly, the code used to perform the analyses presented in this article are available on GitHub (https://github.com/mbernste/spatialcorr-dev).

## Acknowledgements

M.N.B. acknowledges support of a postdoctoral fellowship provided by the Morgridge Institute for Research. C.K., M.N., and Z.N. were supported by GM102756.

## Contributions

M.N.B. and C.K. designed the research project and wrote the first version of the manuscript. M.N. proposed the initial idea to use a log-likelihood ratio for SpatialCorr’s statistical test; and M.N.B. led the model development and validation. A.P. provided domain expertise for the squamous cell carcinoma study. M.N.B. wrote the SpatialCorr package. Z.N., C.M., and J.B. contributed to the code used to test SpatialCorr. All authors contributed to the conceptual development of the paper and to the writing.

## Notes

### Competing Interest Statement

The authors have declared no competing interest.

## References

Andersson, A., and Lundeberg, J. (2021). sepal: identifying transcript profiles with spatial patterns by diffusion-based modeling. Bioinformatics 37, 2644–2650.

Benjamini, Y., and Hochberg, Y. (1995). Controlling the False Discovery Rate: A Practical and Powerful Approach to Multiple Testing. Ournal of the Royal Statistical Society 57, 289–300.

Besag, J., and Clifford, P. (1991). Sequential Monte Carlo p-Values. Biometrika 78, 301–304.

Blondel, V.D., Guillaume, J.-L., Lambiotte, R., and Lefebvre, E. (2008). Fast unfolding of communities in large networks. J Statistical Mech Theory Exp 2008, P10008.

Brown, J., Ni, Z., Mohanty, C., Bacher, R., and Kendziorski, C. (2021). Normalization by distributional resampling of high throughput single-cell RNA-sequencing data. Bioinformatics btab450-.

Cheung, P., Schaffert, S., Chang, S.E., Dvorak, M., Donato, M., Macaubas, C., Foecke, M.H., Li, T.-M., Zhang, L., Coan, J.P., et al. (2021). Repression of CTSG, ELANE and PRTN3-mediated histone H3 proteolytic cleavage promotes monocyte-to-macrophage differentiation. Nat Immunol 22, 711–722.

Demaria, O., Cornen, S., Daëron, M., Morel, Y., Medzhitov, R., and Vivier, E. (2019). Harnessing innate immunity in cancer therapy. Nature 574, 45–56.

Dotto, G.P., and Rustgi, A.K. (2016). Squamous Cell Cancers: A Unified Perspective on Biology and Genetics. Cancer Cell 29, 622–637.

Edsgärd, D., Johnsson, P., and Sandberg, R. (2018). Identification of spatial expression trends in single-cell gene expression data. Nat Methods 15, 339–342.

Ferreira, R.M., Sabo, A.R., Winfree, S., Collins, K.S., Janosevic, D., Gulbronson, C.J., Cheng, Y.-H., Casbon, L., Barwinska, D., Ferkowicz, M.J., et al. (2021). Integration of spatial and single-cell transcriptomics localizes epithelial cell–immune cross-talk in kidney injury. Jci Insight 6, e147703.

Foster, D.S., Januszyk, M., Yost, K.E., Chinta, M.S., Gulati, G.S., Nguyen, A.T., Burcham, A.R., Salhotra, A., Ransom, R.C., Henn, D., et al. (2021). Integrated spatial multiomics reveals fibroblast fate during tissue repair. Proc National Acad Sci 118, e2110025118.

Ghazanfar, S., Lin, Y., Su, X., Lin, D.M., Patrick, E., Han, Z.G., Marioni, J.C., and Yang, J.Y.H. (2020). Investigating higher order interactions in single cell data with scHOT. Nat Methods 17, 799–806.

Haghverdi, L., Büttner, M., Wolf, F.A., Buettner, F., and Theis, F.J. (2016). Diffusion pseudotime robustly reconstructs lineage branching. Nat Methods 13, 845–848.

Han, W., Hu, C., Fan, Z.-J., and Shen, G.-L. (2021). Transcript levels of keratin 1/5/6/14/15/16/17 as potential prognostic indicators in melanoma patients. Sci Rep-Uk 11, 1023.

Harris, C.R., Millman, K.J., Walt, S.J. van der, Gommers, R., Virtanen, P., Cournapeau, D., Wieser, E., Taylor, J., Berg, S., Smith, N.J., et al. (2020). Array programming with NumPy. Nature 585, 357–362.

Hashimshony, T., Feder, M., Levin, M., Hall, B.K., and Yanai, I. (2015). Spatiotemporal transcriptomics reveals the evolutionary history of the endoderm germ layer. Nature 519, 219–222.

Hoff, P.D., and Niu, X. (2012). A Covariance Regression Model. Statistica Sinica 22, 729–753.

Huang, W.-C., Jang, T.-H., Tung, S.-L., Yen, T.-C., Chan, S.-H., and Wang, L.-H. (2019). A novel miR-365-3p/EHF/keratin 16 axis promotes oral squamous cell carcinoma metastasis, cancer stemness and drug resistance via enhancing β5-integrin/c-met signaling pathway. J Exp Clin Cancer Res Cr 38, 89.

Huisman, S.M.H., Lew, B. van, Mahfouz, A., Pezzotti, N., Höllt, T., Michielsen, L., Vilanova, A., Reinders, M.J.T., and Lelieveldt, B.P.F. (2017). BrainScope: interactive visual exploration of the spatial and temporal human brain transcriptome. Nucleic Acids Res 45, e83–e83.

Hunter, M.V., Moncada, R., Weiss, J.M., Yanai, I., and White, R.M. (2021). Spatially resolved transcriptomics reveals the architecture of the tumor-microenvironment interface. Nat Commun 12, 6278.

Inouye, D.I., Yang, E., Allen, G.I., and Ravikumar, P. (2017). A review of multivariate distributions for count data derived from the Poisson distribution. Wiley Interdiscip Rev Comput Statistics 9.

Ji, A.L., Rubin, A.J., Thrane, K., Jiang, S., Reynolds, D.L., Meyers, R.M., Guo, M.G., George, B.M., Mollbrink, A., Bergenstråhle, J., et al. (2020). Multimodal Analysis of Composition and Spatial Architecture in Human Squamous Cell Carcinoma. Cell 182, 497–514.e22.

Karantza, V. (2011). Keratins in health and cancer: more than mere epithelial cell markers. Oncogene 30, 127–138.

Li, J., Luo, H., Wang, R., Lang, J., Zhu, S., Zhang, Z., Fang, J., Qu, K., Lin, Y., Long, H., et al. (2016). Systematic Reconstruction of Molecular Cascades Regulating GP Development Using Single-Cell RNA-Seq. Cell Reports 15, 1467–1480.

Li, Q., Zhang, M., Xie, Y., and Xiao, G. (2021). Bayesian modeling of spatial molecular profiling data via Gaussian process. Bioinformatics.

Liu, Z., Yu, S., Ye, S., Shen, Z., Gao, L., Han, Z., Zhang, P., Luo, F., Chen, S., and Kang, M. (2020). Keratin 17 activates AKT signalling and induces epithelial-mesenchymal transition in oesophageal squamous cell carcinoma. J Proteomics 211, 103557.

Maier, B., Leader, A.M., Chen, S.T., Tung, N., Chang, C., LeBerichel, J., Chudnovskiy, A., Maskey, S., Walker, L., Finnigan, J.P., et al. (2020). A conserved dendritic-cell regulatory program limits antitumour immunity. Nature 580, 257–262.

Maniatis, S., Äijö, T., Vickovic, S., Braine, C., Kang, K., Mollbrink, A., Fagegaltier, D., Andrusivová, Ž., Saarenpää, S., Saiz-Castro, G., et al. (2019). Spatiotemporal dynamics of molecular pathology in amyotrophic lateral sclerosis. Science 364, 89–93.

Maynard, K.R., Collado-Torres, L., Weber, L.M., Uytingco, C., Barry, B.K., Williams, S.R., Catallini, J.L., Tran, M.N., Besich, Z., Tippani, M., et al. (2021). Transcriptome-scale spatial gene expression in the human dorsolateral prefrontal cortex. Nat Neurosci 24, 425–436.

Moll, R., Divo, M., and Langbein, L. (2008). The human keratins: biology and pathology. Histochem Cell Biol 129, 705–733.

Moncada, R., Barkley, D., Wagner, F., Chiodin, M., Devlin, J.C., Baron, M., Hajdu, C.H., Simeone, D.M., and Yanai, I. (2020). Integrating microarray-based spatial transcriptomics and single-cell RNA-seq reveals tissue architecture in pancreatic ductal adenocarcinomas. Nat Biotechnol 38, 333–342.

Nair, R.R., Hsu, J., Jacob, J.T., Pineda, C.M., Hobbs, R.P., and Coulombe, P.A. (2021). A role for keratin 17 during DNA damage response and tumor initiation. Proceedings of the National Academy of Sciences 118, e2020150118.

Paramio, J.M., Casanova, M.L., Segrelles, C., Mittnacht, S., Lane, E.B., and Jorcano, J.L. (1999). Modulation of Cell Proliferation by Cytokeratins K10 and K16. Molecular and Cell Biology 19, 3086–3094.

Paul, F., Arkin, Y., Giladi, A., Jaitin, D.A., Kenigsberg, E., Keren-Shaul, H., Winter, D., Lara-Astiaso, D., Gury, M., Weiner, A., et al. (2015). Transcriptional Heterogeneity and Lineage Commitment in Myeloid Progenitors. Cell 163, 1663–1677.

Quigley, D.A., Kandyba, E., Huang, P., Halliwill, K.D., Sjölund, J., Pelorosso, F., Wong, C.E., Hirst, G.L., Wu, D., Delrosario, R., et al. (2016). Gene Expression Architecture of Mouse Dorsal and Tail Skin Reveals Functional Differences in Inflammation and Cancer. Cell Reports 16, 1153–1165.

Rowe, D.E., Carroll, R.J., and Jr., C.L.D. (1992). Prognostic factors for local recurrence, metastasis, and survival rates in squamous cell carcinoma of the skin, ear, and lip. Journal of the American Academy of Dermatology 26, 976–990.

Rubió-Casadevall, J., Hernandez-Pujol, A.M., Ferreira-Santos, M.C., Morey-Esteve, G., Vilardell, L., Osca-Gelis, G., Vilar-Coromina, N., and Marcos-Gragera, R. (2016). Trends in incidence and survival analysis in non-melanoma skin cancer from 1994 to 2012 in Girona, Spain: A population-based study. Cancer Epidemiol 45, 6–10.

Srivatsan, S.R., Regier, M.C., Barkan, E., Franks, J.M., Packer, J.S., Grosjean, P., Duran, M., Saxton, S., Ladd, J.J., Spielmann, M., et al. (2021). Embryo-scale, single-cell spatial transcriptomics. Science 373, 111–117.

Sun, S., Zhu, J., and Zhou, X. (2020). Statistical analysis of spatial expression patterns for spatially resolved transcriptomic studies. Nature Methods 17, 193–200.

Svensson, V., Teichmann, S.A., and Stegle, O. (2018). SpatialDE: identification of spatially variable genes. Nat Methods 15, 343–346.

Toivola, D.M., Boor, P., Alam, C., and Strnad, P. (2015). Keratins in health and disease. Curr Opin Cell Biol 32, 73–81.

Watson, M.J., Vignali, P.D.A., Mullett, S.J., Overacre-Delgoffe, A.E., Peralta, R.M., Grebinoski, S., Menk, A.V., Rittenhouse, N.L., DePeaux, K., Whetstone, R.D., et al. (2021). Metabolic support of tumor-infiltrating regulatory T cells by lactic acid. Nature 591, 645–651.

Wolf, F.A., Angerer, P., and Theis, F.J. (2018). SCANPY: large-scale single-cell gene expression data analysis. Genome Biol 19, 15.

Yin, J., Geng, Z., Li, R., and Wang, H. (2010). Nonparametric Covariance Model. Stat Sinica 20, 469–479.

Zhao, E., Stone, M.R., Ren, X., Guenthoer, J., Smythe, K.S., Pulliam, T., Williams, S.R., Uytingco, C.R., Taylor, S.E.B., Nghiem, P., et al. (2021). Spatial transcriptomics at subspot resolution with BayesSpace. Nat Biotechnol 39, 1375–1384.

