## Supplementary Materials for "SpatialCorr: Identifying Gene Sets with Spatially Varying Correlation Structure"

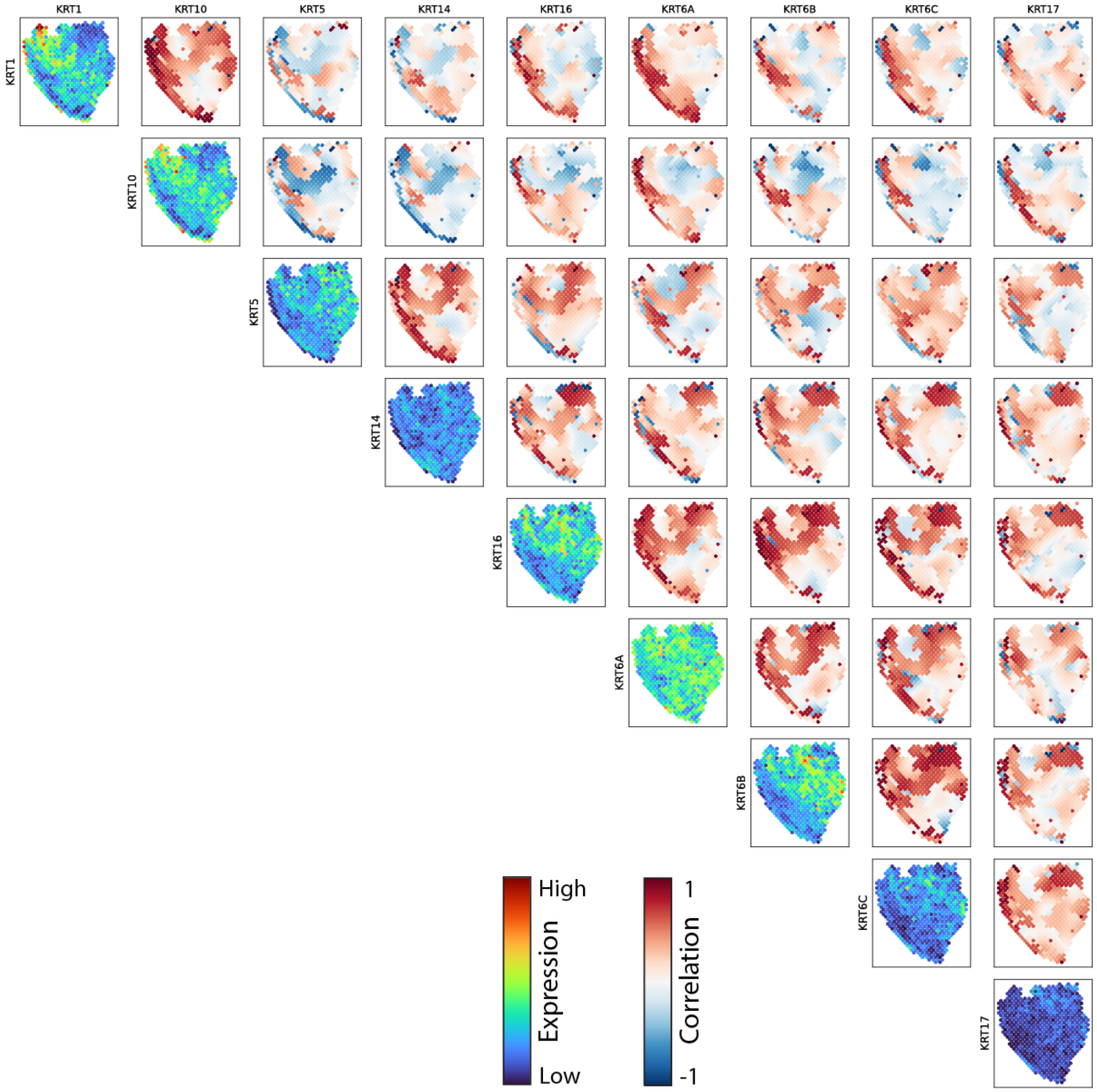


**Supplemental Figure SF1. Estimated correlation between keratin genes in Patient 10.** Off-diagonal heatmaps depict the spot-specific kernel estimates of correlation between all pairs of keratin genes in the "Keratinocyte Differentiation" GO category. Heatmaps along the diagonal depict the spot-specific expression of each gene. No effective-neighbors filter was applied.

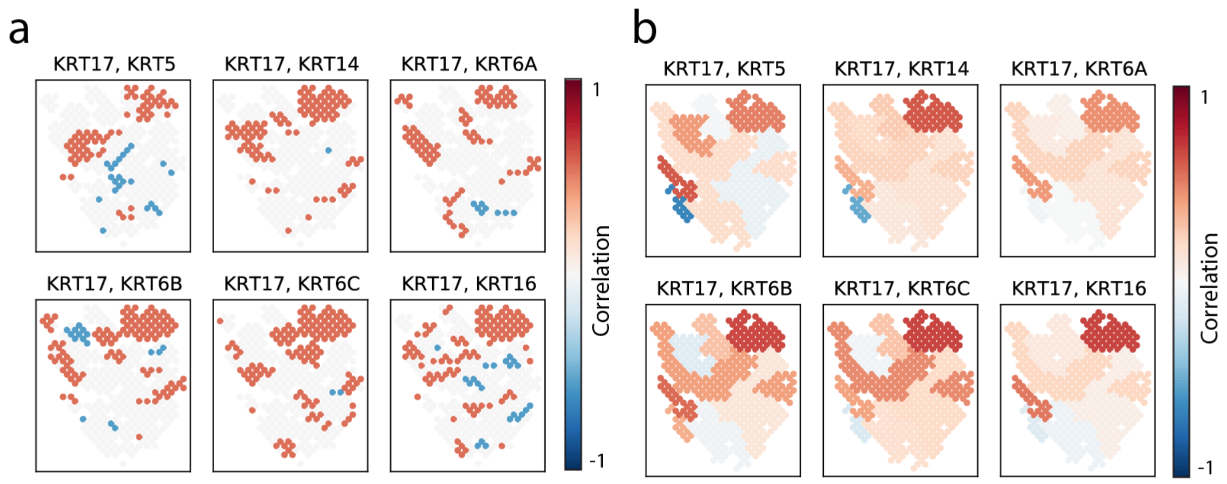


**Supplemental Figure SF2. Example visualizations produced by the SpatialCorr package.** In addition to calculation and visualization of kernel estimates of correlation (as seen in the right panel of Figure 3h), the SpatialCorr package also enables one to calculate and visualize (**a**) approximate confidence intervals around those estimates. A spot is colored red if the 95% confidence interval around the correlation estimate lies above zero. A spot is colored blue if the interval lies below zero. Otherwise, the spot is colored grey. (**b**) SpatialCorr also calculates and visualizes the correlations estimated on a per-region basis using all spots in a region.

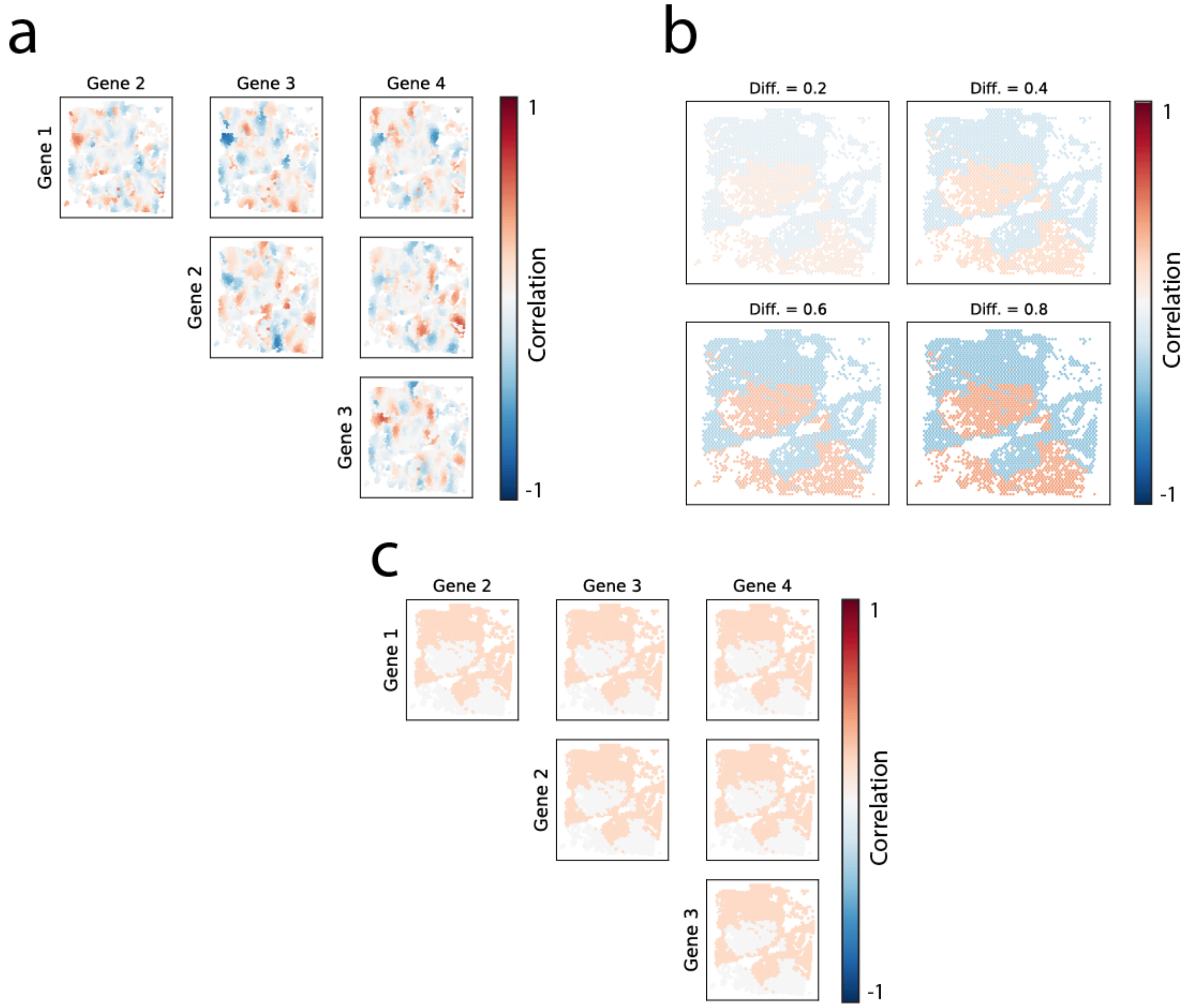


**Supplemental Figure SF3. Example simulated correlation patterns.** Heatmaps showing one instance of the spot-wise latent correlations used to simulate (**a**) four genes in Sim II, (**b**) pairs of genes in Sim III, and (**c**) four genes in Sim IV.

**Table S1: Top five GO terms identified by the WR-test from patient 10**

| GO term | Genes | P-value | Adj. p-value |
|---|---|---|---|
| Keratinocyte differentiation | CSTA, FLG, KRT1, KRT10, KRT14, KRT16, KRT17, KRT5, KRT6A, KRT6B, KRT6C, PERP, S100A7, SFN, SPRR1B | 0.0001 | 0.0103 |
| Epidermis development | FLG, KRT1, KRT10, KRT14, KRT16, KRT17, KRT5, KRT6A, KRT6B, KRT6C, KRTDAP, PERP, S100A7, SFN, SPRR1B | 0.0001 | 0.0103 |
| Epithelial cell differentiation | CSTA, FLG, KRT1, KRT10, KRT14, KRT16, KRT17, KRT5, KRT6A, KRT6B, KRT6C, PERP, S100A7, SFN, SPRR1B | 0.0001 | 0.0103 |
| Regulation of calcium mediated signaling | ATP2B4, CALM1, CALM2, CHP1, CIB1, ERBB3, FKBP1A, GSTO1, HINT1, PRNP, PTBP1, SPPL3 | 0.0019 | 0.1314 |
| Intrinsic apoptotic signaling pathway in response to endoplasmic reticulum stress | ATF4, BAG6, CASP4, CEBPB, ERO1A, ERP29, GRINA, HERPUD1, PARK7, SERINC3, TMBIM6, UBE2K, XBP1 | 0.0021 | 0.1314 |

**Table S2: Top five GO terms identified by the BR-test from patient 10**

| GO term | Genes | P-value | Adj. p-value |
|---|---|---|---|
| Leukocyte migration | ANXA1, ATP1B3, CD44, CD74, COL1A1, COL1A2, CXCL14, MYH9, PPIA, RPS19, S100A14, S100A7, S100A8, S100A9, SDC1 | 0.0001 | 0.031 |
| Myeloid leukocyte migration | ANXA1, CD74, CD81, CD9, CD99, CXADR, HMGB1, IL1RN, RAC1, RPS19, S100A14, S100A7, S100A8, S100A9, WDR1 | 0.0005 | 0.075 |
| Negative regulation of endoplasmic reticulum stress induced intrinsic apoptotic signaling pathway | GRINA, HERPUD1, PARK7, TMBIM6, XBP1 | 0.0062 | 0.6301 |
| Granulocyte migration | ANXA1, CD74, CD99, CXADR, IL1RN, LGALS3, MAPK14, MAPK3, PECAM1, RAC1, S100A14, S100A7, S100A8, S100A9, WDR1 | 0.0090 | 0.6301 |
| Regulation of apoptotic signaling pathway | CD44, CD74, ENO1, GSTP1, HSPB1, RACK1, RPL11, RPL26, RPS3, RPS7, S100A8, S100A9, SFN, TPT1, YWHAZ | 0.0112 | 0.6301 |

**Table S3: Top five GO terms identified by the WR-test from patient 6**

| GO term | Genes | P-value | Adj. p-value |
|---|---|---|---|
| Keratinocyte differentiation | DSP, KRT1, KRT10, KRT14, KRT16, KRT17, KRT5, KRT6A, KRT6B, KRT6C, PERP, PI3, S100A7, SFN, SPRR1B | 0.0001 | 0.002 |
| Epidermis development | FABP5, KRT1, KRT10, KRT14, KRT16, KRT17, KRT5, KRT6A, KRT6B, KRT6C, KRTDAP, PERP, S100A7, SFN, SPRR1B | 0.0001 | 0.002 |
| Cell-cell adhesion | ACTB, ACTG1, ANXA2, CD74, CSTA, DSP, HLA-E, HSPB1, JUP, PERP, PKP1, RPS3, S100A11, S100A8, S100A9 | 0.0001 | 0.002 |
| Tissue migration | ANXA1, CALR, DCN, GRN, HSPB1, JUP, KRT16, KRT2, MYH9, PFN1, RHOA, S100A2, SPARC, TACSTD2, TMSB4X | 0.0001 | 0.002 |
| Myeloid leukocyte migration | ANXA1, CD74, HMGB1, RAC1, RPS19, S100A14, S100A7, S100A8, S100A9 | 0.0001 | 0.002 |

**Table S4: Top five GO terms identified by the BR-test from patient 6**

| GO term | Genes | P-value | Adj. p-value |
|---|---|---|---|
| Apoptotic signaling pathway | CD44, CD74, GSTP1, HSPB1, IFI27, IFI6, PERP, RACK1, RPL11, RPL26, RPS3, S100A8, S100A9, SFN, TPT1 | 0.0001 | 0.0006 |
| Epidermis development | FABP5, KRT1, KRT10, KRT14, KRT16, KRT17, KRT5, KRT6A, KRT6B, KRT6C, KRTDAP, PERP, S100A7, SFN, SPRR1B | 0.0001 | 0.0006 |
| Negative regulation of epithelial cell proliferation | B2M, CAV1, GJA1, SFN, SLURP1, SPARC, STAT1 | 0.0001 | 0.0006 |
| B cell mediated immunity | C1QA, CD74, HLA-E, IGHA1, IGHG1, IGHG3, IGHG4, IGKC, IGLC2 | 0.0001 | 0.0006 |
| B cell activation | BST2, CD74, IGHA1, IGHG1, IGHG3, IGHG4, IGKC, IGLC2, JAK3, LGALS1, MZB1, XBP1 | 0.0001 | 0.0006 |

**Table S5: Top five GO terms identified by the WR-test from patient 4**

| GO term | Genes | P-value | Adj. p-value |
|---|---|---|---|
| Keratinocyte differentiation | CSTA, DSP, KRT1, KRT10, KRT14, KRT16, KRT17, KRT5, KRT6A, KRT6B, KRT6C, PERP, S100A7, SFN, SPRR1B | 0.0001 | 0.0012 |
| Cell cycle | ACTB, CFL1, DYNLL1, EIF4G2, GJA1, HSP90AA1, RACK1, RPL23, RPL24, RPL26, RPS15A, RPS3, RPS6, SFN, TUBA1C | 0.0001 | 0.0012 |
| DNA repair | HMGB1, HMGN1, ISG15, MORF4L1, NPM1, POLR2L, RBX1, RPS27A, RPS3, UBA52, UBB, UBC, UBE2D3, VCP, XRCC6 | 0.0001 | 0.0012 |
| Myeloid leukocyte migration | ANXA1, CD47, CD74, CD81, CD9, HMGB1, LGALS3, MIF, RAC1, RPS19, S100A14, S100A7, S100A8, S100A9, WDR1 | 0.0001 | 0.0012 |
| T cell proliferation | ANXA1, CD24, CD81, CEBPB, CTNNB1, GPNMB, HLA-E, HMGB1, LGALS3, PRDX2, PRKAR1A, PRNP, PYCARD, RPS3, RPS6 | 0.0001 | 0.0012 |

**Table S6: Top five GO terms identified by the BR-test from patient 4**

| GO term | Genes | P-value | Adj. p-value |
|---|---|---|---|
| Keratinocyte proliferation | CDH3, EPPK1, IRF6, KLK8, KRT2, NOTCH2, SDR16C5, SFN, SLURP1, SRSF6, TGM1, TP63, YAP1, ZFP36, ZFP36L1 | 0.0001 | 0.0008 |
| Cell cycle | ACTB, CFL1, DYNLL1, EIF4G2, GJA1, HSP90AA1, RACK1, RPL23, RPL24, RPL26, RPS15A, RPS3, RPS6, SFN, TUBA1C | 0.0001 | 0.0008 |
| Cell matrix adhesion | CD44, CD63, COL17A1, COL3A1, CTNNB1, CTTN, EMP2, ITGA6, ITGB1, JUP, LYPD3, MACF1, RAC1, RHOA, S100A10 | 0.0001 | 0.0008 |
| Response to interferon gamma | ACTG1, ACTR3, B2M, CD44, CDC42, GAPDH, GSN, HLA-DRA, HLA-E, HSP90AB1, IFITM3, MT2A, RPL13A, STAT1, VIM | 0.0001 | 0.0008 |
| Regulation of cell death | GAPDH, GJA1, GSTP1, HSPB1, MT-CO2, PERP, RPL10, RPL26, RPS27A, RPS3A, RPS6, S100A8, S100A9, SFN, TPT1 | 0.0001 | 0.0008 |

**Table S7: Top five GO terms identified by the WR-test from patient 2**

| GO term | Genes | P-value | Adj. p-value |
|---|---|---|---|
| Keratinocyte differentiation | CSTA, JUP, KRT1, KRT10, KRT14, KRT16, KRT17, KRT5, KRT6A, KR T6B, KRT6C, PERP, S100A7, SFN, SPRR1B | 0.0001 | 0.0117 |
| Cell cell adhesion | ACTB, ACTG1, ANXA1, ANXA2, CD74, CSTA, DSP, GNAS, JUP, LG ALS7B, PERP, RPS3, S100A11, S10 0A8, S100A9 | 0.0001 | 0.0117 |
| Regulation of epidermis development | AQP3, CDH3, CDSN, CTNNB1, CT SK, HES1, KRT10, KRT17, KRT2, RUNX1, SERPINB13, SFN, SPINK5 , TP63, ZFP36L1 | 0.0001 | 0.0117 |
| Epidermis development | CSTA, FABP5, KRT1, KRT10, KRT 14, KRT16, KRT17, KRT5, KRT6A, KRT6B, KRT6C, PERP, S100A7, SF N, SPRR1B | 0.0001 | 0.0117 |
| Epithelial cell differentiation | CSTA, JUP, KRT1, KRT10, KRT14, KRT16, KRT17, KRT5, KRT6A, KR T6B, KRT6C, PERP, S100A7, SFN, SPRR1B | 0.0001 | 0.0117 |

**Table S8: Top five GO terms identified by the BR-test from patient 2**

| GO term | Genes | P-value | Adj. p-value |
|---|---|---|---|
| Keratinocyte differentiation | CSTA, JUP, KRT1, KRT10, KRT14, KRT16, KRT17, KRT5, KRT6A, KRT6B, KRT6C, PERP, S100A7, SFN, SPRR1B | 0.0001 | 0.0031 |
| Positive regulation of calcium ion transmembrane transport | ABL1, ATP1B1, CALM1, CALM2, CALM3, CXCL10, CXCL9, G6PD, GSTO1, HSPA2, HTT, SRI, STIM1, SUMO1, THY1 | 0.0001 | 0.0031 |
| Regulation of cell death | CFL1, ENO1, GSTP1, LDHA, PERP, RACK1, RPL11, RPS27A, RPS3, RPS6, S100A8, S100A9, SFN, TPT1, UBA52 | 0.0001 | 0.0031 |
| Positive regulation of cell cell adhesion | ANXA1, CD24, CD44, CD47, CD74, CD81, CDC42, HMGB1, IRAK1, ITGA6, LGALS1, RAC1, RHOA, RPS3, THY1 | 0.0001 | 0.0031 |
| Cell matrix adhesion | ACTN1, CD44, CD63, COL17A1, COL3A1, CTNNB1, FN1, ITGA6, ITGB5, JUP, LYPD3, RAC1, RHOA, S100A10, THY1 | 0.0001 | 0.0031 |

**Supplementary Note S1. SpatialCorr avoids false positives due to abrupt changes in mean expression between regions:** Spurious correlations can be induced by changes in latent factors such as mean expression. The figure below provides a simple example generated by the Poisson-lognormal model in which there exist two tissue regions with large differences in mean expression, but no spatially varying correlation within each region. The differences in mean expression induce spuriously high correlation along the border between the regions, which leads to false identification of spatially varying correlation when not taking into account the differing means between the regions.

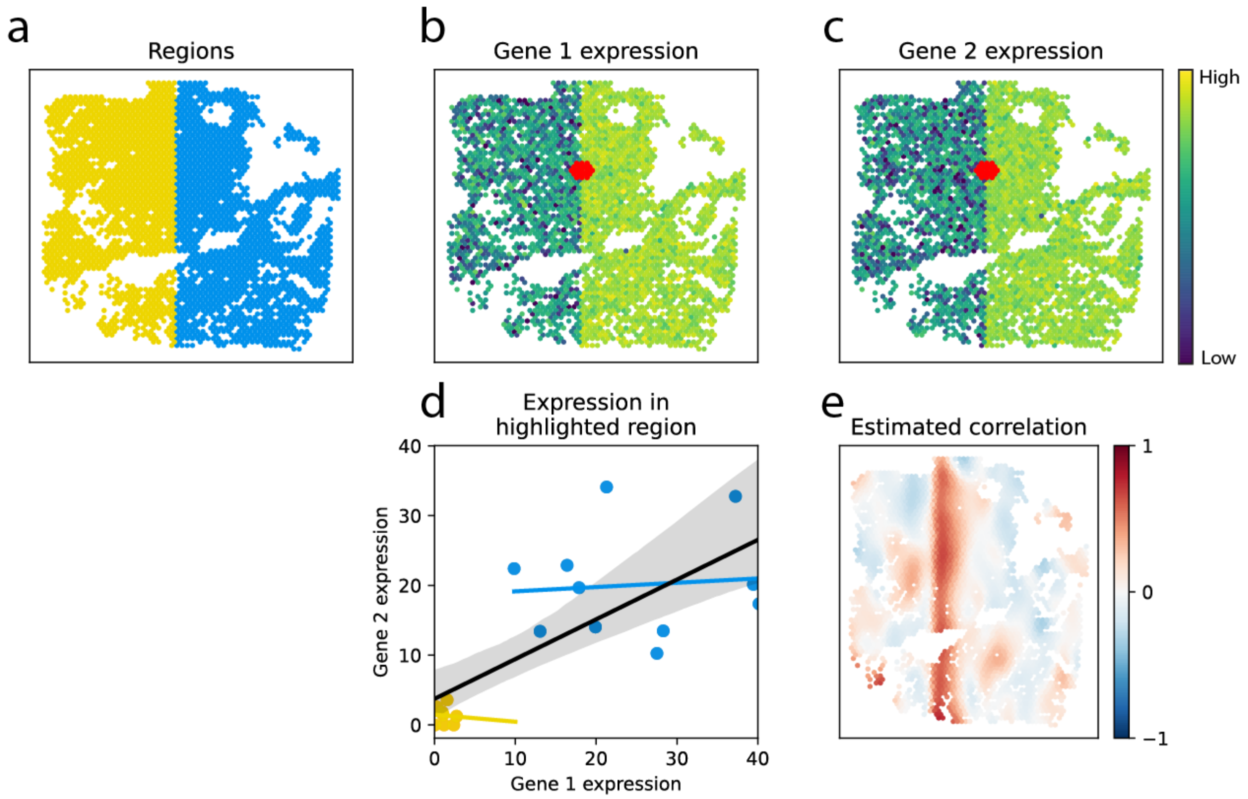


**Supplementary Figure SF3. Example of spurious spatial correlation.** We depict a toy example in which the slide is (**a**) comprised of two regions (yellow and blue) where the two genes' mean expression differs between the two regions (**b**,**c**), but the correlation between these two genes is zero within each region. We highlight a region of spots (red) along the border between the two regions and show a scatterplot of the expression values for the spots within this region (**d**). Points are colored according to the region to which they belong. The yellow line shows the least-squares regression line for points in the yellow region; the blue line shows the regression line for points in the blue region. The slopes of each are near zero which is expected since the genes are not correlated within region. Also shown in solid black is the least-squares regression line fit to points from both regions; 95% error regions are generated from bootstrapping. Here, the genes appear to be correlated due to differences in mean expression. The kernel estimates of the correlation at each spot shows a band of high correlation along the border between the two regions (**e**).

To evaluate the effect of differences in latent mean expression across tissue regions, we considered 200 datasets simulated from the Poisson-lognormal-based model as shown in Figure SF3. SpatialCorr and scHOT were applied to each dataset; the p-value distributions for each approach are shown below. By conditioning on tissue type, SpatialCorr avoids false positives and yields uniformly distributed p-values under the null distribution. In contrast, scHOT, which does not condition on tissue type, produces high rates of false positives. To address this in practice, scHOT restricts analyses to gene pairs showing no change in average expression. Given that most interesting genes will show some change in mean expression across a heterogeneous tissue, limiting the search to non-SV genes is often restrictive.

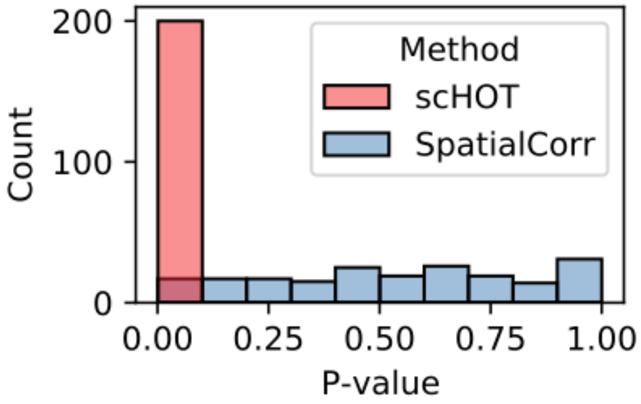


**Supplementary Figure SF4. Comparing SpatialCorr to scHOT on simulated data with differing means between regions.** The distribution of p-values produced by scHOT and SpatialCorr when run on 200 simulated datasets similar to that shown in Figure SF3.

**Supplementary Note S2. Comparison of correlation in simulated data versus experimental data):** We sought to generate latent correlation patterns (such as those shown in Figure 2c) that match the latent correlations between genes in experimental data. This task is challenging because the known latent correlations used as input to the Poisson-lognormal simulation model are not directly comparable to the estimated correlations from the experimental data. In short, the fewer the UMI counts, the greater the discrepancy between the latent and observed correlation (Wang et al., 2020). Unfortunately, this issue is not circumvented using normalization methods that attempt to infer the latent expression values in the experimental data, such as Dino (Brown et al., 2021) and SAVER (Huang et al., 2018), due to the uncertainty in estimates of the latent correlation values.

To assess how well our simulated correlation patterns match those found in experimental data, we performed an empirical comparison of the spot-wise correlations estimated from the normalized counts (via the Gaussian kernel) between the simulated and experimental data.

Specifically, we performed the following analysis:

1. For each level of latent correlation (i.e., weak, moderately weak, moderately strong, or strong), we generate 10 simulated datasets using experimental data for the Rank 50 gene to seed the simulation and normalize each simulated dataset.
2. For each simulated dataset, we estimate the spot-wise correlations using Gaussian kernel estimation. We then compute the variance of these spot-wise correlation estimates across the slide.
3. For all pairs of genes between ranks 40-60, we estimate the spot-wise correlations using the experimental data, and for each pair, compute the variance of these spot-wise correlation estimates.

We found the estimated correlations calculated from the simulated data are similar to those observed in the experimental data at each of the four expression levels. The figure below shows that the distributions of spot-wise correlations from simulated data are similar to those found in experimental data.

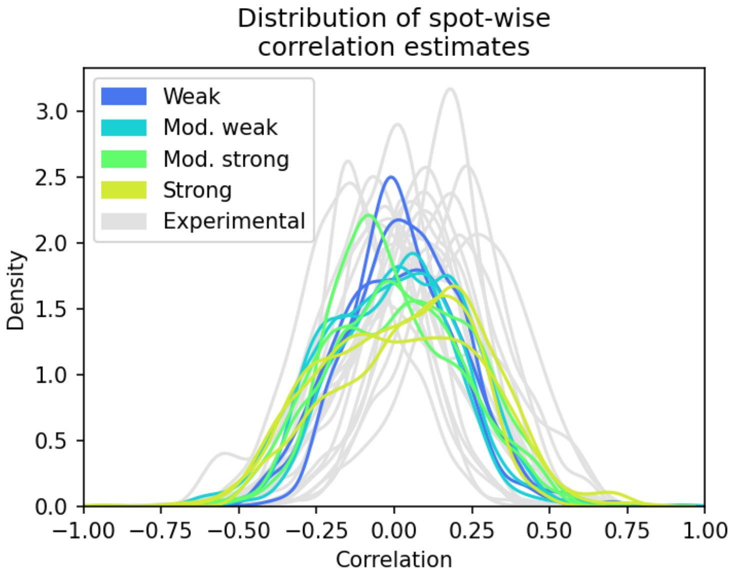


**Supplementary Figure SF5. Distribution of spot-wise correlations in simulated and experimental data.** The kernel density estimated distributions of the spot-wise correlations in simulated (colorful) and experimental (grey) datasets. Specifically, we sample 10 random pairs of genes whose expression falls within ranks 40-60 from the experimental data, estimate each-spot's correlation between each pair using Guassian kernel estimation, and for each pair, plot the distribution of those per-spot correlations across the slide. For each latent correlation level, we sample three simulated datasets and repeat this procedure.

To more rigorously assess how the variability of simulated spot-wise correlation estimates compare to those in experimental data, below we plot the distribution of the variances of the spot-wise correlation values for all gene-pairs assessed in the experimental data. For each latent correlation level, the average variances of the correlation estimates across all simulated datasets generated with that latent correlation level level is shown (average taken over 10 replicates). As seen below, we found that the average variance for the highest latent correlation level level lay within the tail of the distribution of the variances calculated from the experimental data, suggesting that the highest latent correlation level used in the simulations corresponds to a typical strongly correlated gene-pair in real data.

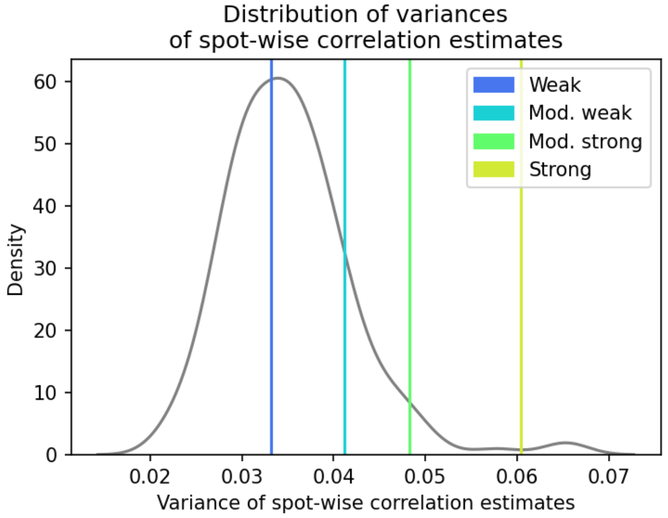


**Supplementary Figure SF6. Variance of spot-wise correlations in simulated and experimental data.** For each gene pair within ranks 40-60 in the experimental data, we estimate each spot's correlation using Gaussian kernel estimation, and for each pair, compute the variance of those spot-wise correlation estimates across the slide. Shown is the distribution of these empirical variances across all gene pairs (grey). For each latent correlation level level, we repeat this procedure, calculate the average variance (taken over 10 replicates), and plot these average values above (colorful vertical lines).

### References


Brown, J., Ni, Z., Mohanty, C., Bacher, R., and Kendziorski, C. (2021). Normalization by distributional resampling of high throughput single-cell RNA-sequencing data. Bioinformatics btab450-.

Huang, M., Wang, J., Torre, E., Dueck, H., Shaffer, S., Bonasio, R., Murray, J.I., Raj, A., Li, M., and Zhang, N.R. (2018). SAVER: Gene expression recovery for single-cell RNA sequencing. Nat Methods *15*, 539–542.

Wang, Y., Hicks, S.C., and Hansen, K.D. (2020). Co-expression analysis is biased by a mean-correlation relationship. Biorxiv 2020.02.13.944777.
